# Minor Cannabinoids CBD, CBG, CBN and CBC differentially modulate sensory neuron activation

**DOI:** 10.1101/2025.10.02.680148

**Authors:** Katalin Rabl, Larry Gruenke, Ashwin Banfal, Helge Eilers, Judith Hellman, Mark A Schumacher

## Abstract

The use of minor cannabinoids has been advanced, in part, by the idea of providing relief from pain and inflammation without the burden of unwanted psychogenic effects associated with Δ^9^THC. In this regard, investigators have focused on the effects of minor cannabinoid activation / desensitization of peripheral sensory neurons on nociceptive signaling and/or peripheral inflammation. With a focus on peripheral nociception, four common minor cannabinoids: cannabidiol (CBD), cannabigerol (CBG), cannabinol (CBN) and cannabichromene (CBC) were studied in primary cultures of mouse Dorsal Root Ganglion (DRG) neurons. We queried if calcium responses induced by the four cannabinoids differed in potency of activation, neuronal size preference, and dose-response relationships. Additionally, we determined the dependence of CBD and CBN on key channel-receptors that are known to mediate pain and/or antinociception. Individually, CBD, CBG and CBC directed greater response magnitudes when compared to CBN. All four minor cannabinoids activated overlapping but distinct size populations of sensory neurons. CBD and CBG activated the widest range of DRG neuron sizes (smaller-larger) overlapping with smaller capsaicin-sensitive neurons. In contrast, CBN and CBC activated predominantly larger sensory neurons. CBD diverged from other minor cannabinoids in directing a linear dose-response profile whereas CBG and CBC directed sigmoidal dose-response profiles and CBN activated DRG neurons with an inverted U-shaped dose-response relationship. CBD-induced activation of DRG neurons was dependent on co-expression of the nociceptive channel TRPV1 plus cannabinoid receptor 1 (CB_1_R), whereas CBN-induced activation was independent of TRPV1. Overall, we observed that minor cannabinoids CBD, CBG, CBN and CBC differed in their activation of DRG neurons and directed unique activation properties across a diverse population of sensory neurons. Such differences underly the hypothesis that a combination (entourage) of complimentary minor cannabinoids can direct synergistic antinociceptive activity.

## Introduction

Minor cannabinoids, a diverse set of diterpene-related molecules derived from the plant *Cannabis sativa*, are purported to have anti-inflammatory and analgesic properties [1]. When compared with Δ^9^THC, the major bioactive cannabinoid in *C. sativa*, minor cannabinoids such as cannabidiol (CBD) have a lower incidence or absence of unwanted central nervous system side effects [2]. Despite the substantial increase in public use of certain minor cannabinoids for the treatment of a wide range of conditions including pain, our understanding of their mechanisms of analgesic action is very limited.

The sensation of pain principally arises from the activation of peripheral nociceptors - specialized primary afferent sensory neurons dedicated to the detection of actual or impending tissue injury [3, 4]. Sensory neurons are functionally distinguished by their unique properties of activation, typically requiring noxious thermal, chemical and/or mechanical stimuli of sufficient magnitude to activate high threshold receptor / channels expressed on nociceptor terminals [5]. Using primary cultures of rodent Dorsal Root Ganglion (DRG) neurons, we and others have modeled properties of nociceptor activation *in vitro* [6-8]. While CBD has been shown to activate and desensitize detectors of noxious stimuli such as the transient receptor potential cation channel subfamily Vanilloid member 1 (TRPV1), comparative studies of minor cannabinoid activation have often been limited to the study of receptor / channels in non-neuronal cell lines (HEK) rather than in primary cultures of sensory neurons [9, 10].

We hypothesized that CBD, CBG, CBN and CBC would differentially activate DRG neurons. Accordingly, we compared the activation properties of these four minor cannabinoids in cultured mouse DRG neurons by quantifying calcium imaging of their relative potency, neuronal size preference for activation and dose-response profiles. Importantly, we examined the divergent activation profiles of CBD and CBN and their dependence on TRPV1 and the cannabinoid receptor subtype -1 (CB_1_R).

## Methods

### Animals

Male and female mice (8-12 weeks) used in this study included: C57Bl/6 (BL6), Wild type (Wt) versus mice with total TRPV1 knockout (TRPV1^-/-^) (Jackson Laboratory, Bar Harbor, ME) [11]; Mice with CNRO (CB_1_R) conditional knockdown in the peripheral nervous system and in sensory neurons were generated by breeding CRNO Flox mice with Advillin Cre [12] or Nav1.8 Cre mice (Jackson Laboratory), respectively. CNRO Flox mice were provided by Josephine Egan MD, NIH [13, 14]. Mice were genotyped, bred and housed within the UCSF laboratory animal resource center (LARC) in a climate-controlled room on a 12-hours light/dark cycle. Genotyping was performed using gDNA from mouse ear clippings processed with the Extract-N-Amp Tissue Direct kit (Sigma-Aldrich). Laboratory diet was freely available to mice ad libitum. Experimental protocols were approved by the University of California, San Francisco, Institutional Animal Care and Use Committee (IACUC).

### Dorsal Root Ganglion (DRG) neuronal cultures

Lumbar DRGs were harvested from adult male and female mice (2 mice for each experimental day) and sensory neurons were cultured as previously described [15].

### Minor Cannabinoid Stock Solutions

Minor cannabinoid stock solutions were prepared from crystalline solid (except for CBC which was sold as a 1mg/ml solution in methanol) (Cayman Chemical) in either a 1:1 solution of 100% ethanol:distilled water (cannabidiol - CBD) or dissolved in pure 100% ethanol: cannabigerol (CBG), cannabinol (CBN) and cannabichromene (CBC) to a final target concentration of 2 mg/ml and stored in glass vials at -20°C. In the case of CBC, methanol was first evaporated using a nitrogen source before resuspending the compound in 100% ethanol.

### HPLC Analysis of Minor Cannabinoids

Analytical chromatography was carried out on an Agilent 1100 HPLC using a 4.6mm × 150mm Agilent Eclipse Plus C18 column (3.5 micron) at a flow rate of 1.2 ml/min with UV detection at 210 nm. Elution (Mobile Phase A vs Mobile Phase B) was optimized for each application: Mobile Phase A = 0.1% TFA in 9:1 water: acetonitrile; Mobile Phase B = 0.1% TFA in 1:9 water:acetonitrile. For analysis of CBD or CBG content in stock solutions, elution was isocratic with 10% Mobile Phase A; 90% Mobile Phase B. For analysis of CBN or CBC content, elution was isocratic with 5% Mobile Phase A; 95% Mobile Phase B. Cannabinoid purity (**Fig. S1**) was assessed using a gradient elution from 40% 0.1% aqueous TFA to 100% aqueous 0.1% TFA vs Mobile Phase B over a period of 8 min then held at 100% 0.1% TFA for an additional 6 min.

### Preparation and Validation of Minor Cannabinoid Aqueous Solutions

Minor cannabinoids from stocks in ethanol were diluted into aqueous calcium imaging (assay) buffer (Hank’s buffered salt solution containing calcium and magnesium with 20mM HEPES, pH 7.4), in glass with minimal vortexing and no sonication. To minimize time- and concentration-dependent loss of cannabinoids prior to injection into the calcium imaging chamber, the desired cannabinoid solution was transferred to a glass Luer Lock syringe installed on a syringe pump and connected to PEEK tubing (OD: 1.59mm, ID 1.016 mm, Trajan Scientific and Medical), allowing cannabinoids to be directly injected into the imaging bath without contact with other plastics. To mitigate an expected time-dependent decrease in the targeted cannabinoid concentration, we expedited cannabinoid experiments and frequently exchanged freshly prepared aqueous cannabinoid solutions in the glass syringe reservoir and PEEK tubing.

### Calcium Imaging and Microscopy

Measurements of cultured lumbar DRG neuron intracellular calcium [Ca^2+^]_i_ and cell size were performed by preloading neurons previously plated on coverslips with 5 μM cell permeant Fluo-4 AM (Invitrogen, Waltham, MA) calcium-sensitive dye (488/520 nm) with Pluronic acid F-127 (Invitrogen) for 45 minutes at 37°C, 5% CO_2_. DRG neurons were visualized through a Zeiss Axiovert microscope (10x) equipped with an Axiocam camera and quantified using Zen Pro software (Carl Zeiss, Jena, Germany) with a Camera ROI: 20-75 cells. Neurons were perfused with Hank buffered salt solution with calcium and magnesium, supplemented with 20 mM HEPES and 1% penicillin–streptomycin, at 2 mL/minute via a valve-controlled system at room temperature (22°C). Responsive neurons are defined as those with an increase of [Ca^2+^]_i_ ≥ 10% from baseline values. Neurons that did not respond to KCl were excluded.

### Reagents

CBD, CBG, CBN, CBC, CB1R antagonist (Rimonabant - SR141716) (Cayman Chemical), Capsaicin (Sigma-Aldrich), TRPV1 antagonist (SB705498) (Apex Bio).

### Statistical Analysis

When applicable, determination of differences between group means (+/-SEM) was performed by ANOVA (eg. comparisons of CBD, CBG, CBN and CBC - induced mean dF/F% peak DRG neuron calcium responses at 50 μM) or two-way unpaired t-tests (eg. mean CBN-induced peak DRG neuron calcium response dF/F% between Wt versus CB_1_R cKD. A *P* value of < 0.05 was considered statistically significant. Statistical analysis was performed by Prism (GraphPad) including use of GraphPad’s non-linear curve fitting for dose-response data, “*Find ECanything*”, best fit analysis (CBD, CBG, CBC) and a second-order polynomial (quadratic) equation for CBN (GraphPad). Figures were generated using Adobe Illustrator.

## Results

Prior studies have primarily focused on the action of minor cannabinoids on individual nociceptive channels / receptors expressed in heterologous expression systems such as HEK cells [9, 10]. Paradoxically, the antinociceptive effects ascribed to these minor cannabinoids have been linked, in part, to their ability to activate then desensitize and/or block pain transducing ion channels in sensory neurons [16]. While CBD, CBG, CBN and CBC have been reported to activate rodent primary sensory neurons including those with nociceptive properties, their comparative potencies, size-dependent activation, dose-response relationships and dependence on TRPV1/CB_1_R have not been fully described [16].

Following validation of minor cannabinoid integrity and concentrations in stock and aqueous bath solutions, we applied calcium imaging techniques capable of surveying populations of cultured DRG neurons to study CBD, CBG, CBN, CBC - induced response relationships as quantified by dF/F% in [Ca^2+^]_i_ and cell size. Additionally, we applied a combination of genetic and pharmacologic strategies to investigate potential differences between CBD and CBN receptor-mediated activation of sensory neurons.

### Validation of minor cannabinoid concentrations in stock and aqueous bath solutions

All cannabinoids obtained were analyzed for purity and stability by HPLC with UV detection at 210 nm and found to be 98-100% pure. Stock solutions were stable at -20°C for up to 8 months and changes in the concentration of the solutions were calculated from the peak areas and impurities were reported as percent of the total peak areas in the chromatogram (**Fig. S1)**. Preparation of aqueous solutions of minor cannabinoids were problematic given their hydrophobic properties and binding to plastics and plastic tubing. Following methods developed and utilized as described (**Fig. S2A-C**), HPLC analysis of aqueous solutions containing cannabinoids demonstrated that they were within 10% of their targeted concentration utilizing a glass syringe reservoir and calcium imaging bath solution delivery via PEEK tubing.

### CBD, CBG, CBN and CBC differ in DRG neuron response magnitudes at 50μM

We observed consistent activation of cultured DRG neurons by CBD, CBG, CBN and CBC at 50μM (**Fig. 1A-D)**. This is consistent with the concentration range (30 -100 μM) that is reported by others to activate cultured DRG neurons by CBD and CBG [16], and/or approximated the EC_50_ for TRPV1 channel activation by CBD (30 μM) and CBN (36 μM) expressed in HEK cells [10]. As quantified by a maximal increase in fluorescence (dF/F%), we observed [Ca^2+^]_i_ response magnitudes of CBD = 70.8 +/-5.9, CBG = 104 +/-11.8, CBN = 33.8 +/-4.1 and CBC = 85.5 +/-5.5% (**Fig. 1E**). Overall, individual response magnitudes of CBD (** P < 0.006), CBG and CBC (**** P <0.0001) were greater than those directed by CBN.

**Fig. 1.**
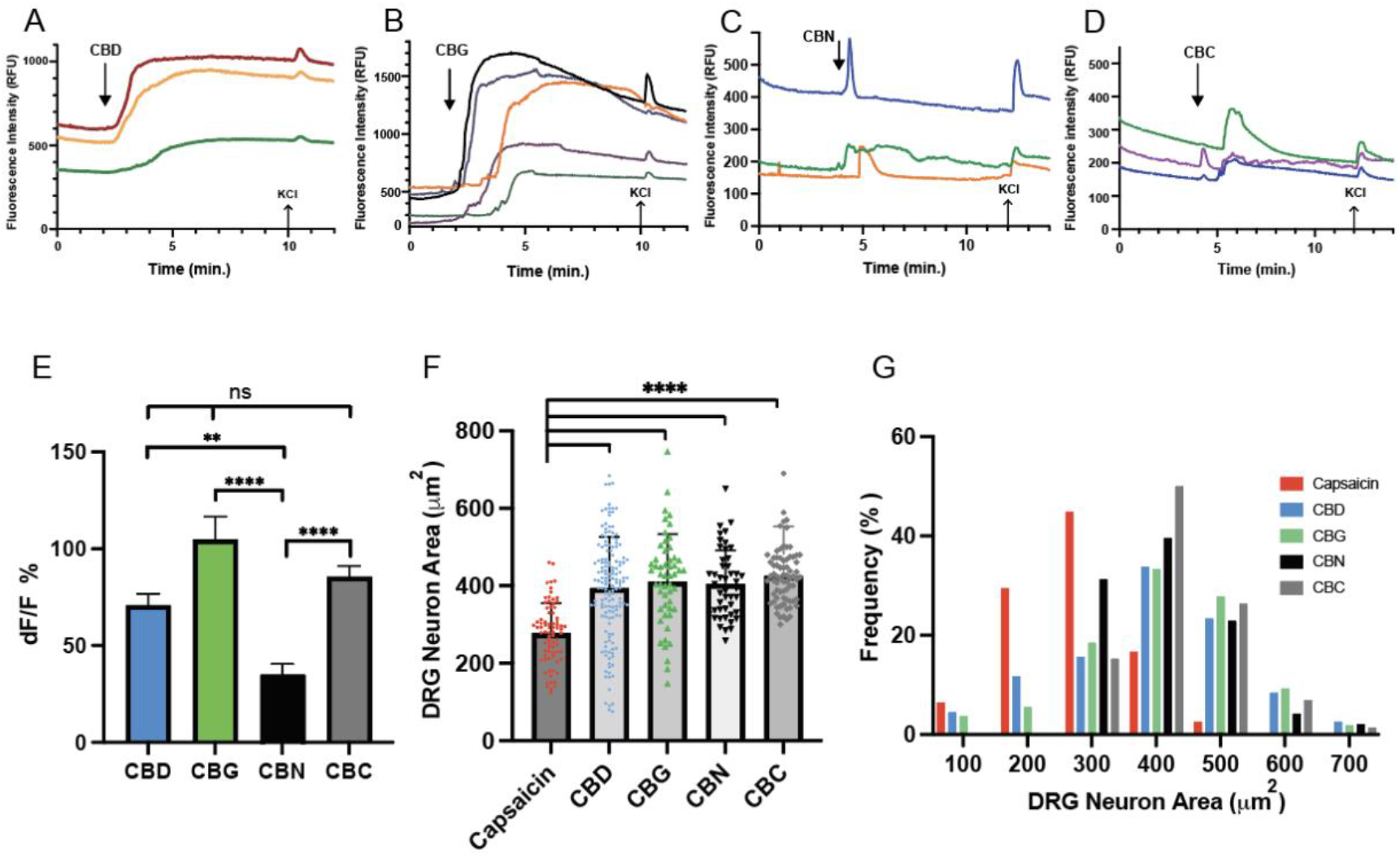
Minor cannabinoids differ in their response magnitude and size-dependent frequency of responses in DRG neurons. Representative [Ca^2+^]_i_ response recordings of **(A)** CBD – cannabidiol, **(B)** cannabigerol – CBG, **(C)** cannabinol – CBN, and **(D)** cannabichromene – CBC at 50μM for 15 sec (top arrows), followed by KCl (bottom arrows), (**E**) Comparison of CBD, CBG, CBN and CBC (50 μM) – induced [Ca^2+^]_i_ responses in cultured DRG neurons. CBN (n=31) directed the smallest calcium response compared to CBD (n=40), CBG (n=28) and CBC (n=35) (** P < 0.006); (**** P < 0.0001) (ns = not significant), (**F**) Size scatter plots of capsaicin and minor cannabinoid-responsive DRG neurons. Capsaicin activated a smaller size (277 μm^2^) subpopulation of DRG neurons (n=78) as compared with those activated by CBD (394 μm^2^) (n=154), CBG 410 μm^2^ (n=54), CBN 406 μm^2^ (n = 48) and CBC 426 μm^2^ (n= 72) at 50μM (**** P <0.0001). (**G**) Size frequency distribution (%) of capsaicin and cannabinoid-responsive DRG neurons. While the mean neuronal size (μm^2^) did not differ between CBD, CBG, CBN and CBC, the majority of DRG neurons > 400 μm^2^ were activated by CBN and CBC. DRG neurons attaining an increase of dF/F% ≥ 10% from baseline over a 3 min recording interval were considered responders in determining size frequency distribution. Mean values are shown with error bars +/-SEM representing values from ≥ 3 independent trials. (dF/F%) is the percent difference from baseline [Ca^2+^]_i_ of the maximal response measured over a three - minute interval. Differences were determined by ANOVA with Sidak’s corrections for multiple comparison (Graph Pad). Only DRG neurons responsive to KCl at the conclusion of the experiment were included.

### Capsaicin, CBD, CBG, CBN and CBC activate overlapping but distinct size populations of sensory neurons

Minor cannabinoids CBD, CBG, CBN and CBC can directly activate TRPV1 in heterologous cell expression systems [10]. Specifically, CBD is proposed to direct its antinociceptive properties, in part, through the activation / desensitization of TRPV1 – expressing sensory neurons [16]. Given that TRPV1 expression is largely restricted to the small-medium sized DRG neurons under control conditions, we compared the size distribution of DRG neurons activated by CBD, CBG, CBN and CBC to those activated by the TRPV1 agonist, capsaicin [15, 17]. As shown in **Fig. 1F**, capsaicin activated a smaller-size (mean area 277 +/-8.7 μm^2^) subpopulation of DRG neurons that differed from the larger mean areas (μm^2^) of neurons activated by CBD (394 +/-10.6); CBG 410 +/-16.8; CBN 406 +/-12.5; and CBC 426 +/-8.5 (**** P<0.0001). However, capsaicin-responsive neurons primarily overlapped in size with those activated by CBD and CBG (a smaller subpopulation of neurons) (**Fig. 1G**), as compared to CBN and CBC that activated predominantly larger DRG neurons with areas greater than 400 μm^2^. Overall, this suggests that CBD, CBG, CBN and CBC may differ in their properties of sensory neuronal activation that can be further characterized by their dose-dependent activation profiles.

### Minor cannabinoids CBD, CBG, CBN and CBC direct distinct dose-response profiles in DRG neurons

To establish a more detailed comparison of properties of sensory neuronal activation induced by CBD, CBG, CBN and CBC, we conducted dose-response experiments across all four minor cannabinoids and observed complexity within and between the minor cannabinoid dose-response profiles. As shown in **Fig. 2A**, CBD induced dose-dependent increases in DRG neuronal [Ca^2+^]_i_ that followed a linear rather than sigmoidal relationship of its dose-response (dF/F%) profile. In contrast, CBG (**Fig. 2B**) directed a dose-dependent increase in [Ca^2+^]_i_ that more closely modeled a sigmoidal response relationship but with the notable exception that the highest CBG concentration tested (100 μM) directed a smaller response than observed at 25 μM or 50 μM. As shown in **Fig. 2C**, CBN directed the most heterogeneous dose-response profile amongst the minor cannabinoids studied. While CBN directed an increase in [Ca^2+^]_i_ from 0.1 μM to 1 μM, greater CBN concentrations induced smaller responses from 10 μM to 100 μM that most closely modeled an inverted U-shaped curve. Finally, CBC (**Fig. 2D**) directed a dose-response profile that, in part, approximated a sigmoidal relationship but includes features of a biphasic response profile. Together these findings reflect the complexity of minor cannabinoid-induced activation across divergent sensory-neuronal phenotypes [18].

**Fig. 2.**
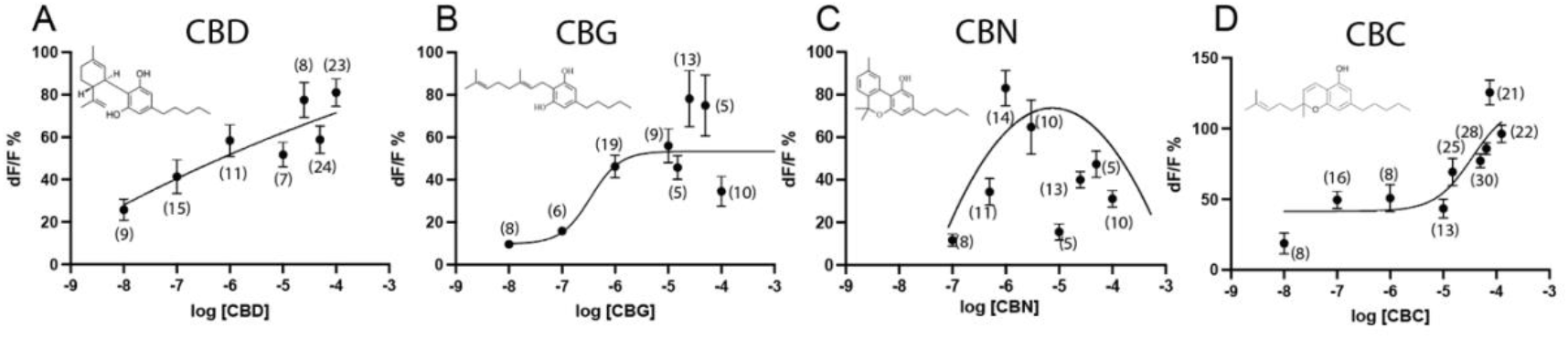
DRG neuron dose-response relationships differ between CBD, CBG, CBN and CBC. Log dose-responses of minor cannabinoids derived from a total population of cultured DRG neurons induced differing profiles. (**A**) CBD directed a linear Log dose-response profile. (**B**) CBG directed a sigmoidal dose-response. (**C**) CBN directed an inverted U-shaped dose-response profile. (**D**) CBC directed a sigmoidal dose-response profile with features of biphasic activation. Log dose-response profiles were plotted based on best curve fits (Prism, GraphPad). Mean values (+/-SEM) are shown for each cannabinoid concentration tested for total DRG neurons (A-D) with (n) independent neurons from ≥ 2 independent trials. dF/F% is the percent difference from baseline [Ca^2+^]_i_ of the maximal response measured over a 3-minute interval. Only DRG neurons responsive to KCl at the conclusion of each experiment were included.

### CBD-induced activation of DRG neurons is dependent on co-expression of TRPV1 and CB_1_R

The phytocannabinoid Δ^9^THC, the synthetic cannabinoid WIN 55212,22 and the endocannabinoid - anandamide are reported to activate TRPV1 expressed alone or co-expressed with CB_1_R [19-21]. However, it remains unclear whether the minor cannabinoids CBD and CBN require TRPV1 and/or CB_1_R to activate DRG neurons. We compared and observed no difference in the calcium response magnitudes of CBD (50 μM) to DRG neurons derived from male Wt (68.5 +/-6.7, n=33) versus TRPV1 KO (58.8 +/-8.6, n=16) mice. Likewise, we studied the response magnitude of CBD in the presence of the CB_1_R antagonist Rimonabant [19], (62.5 +/-6.6, n=9) and also observed no change in the CBD-induced response magnitudes. Unexpectedly, when the combination of DRG neurons derived from TRPV1 KO mice were challenged with CBD in the presence of Rimonabant, no CBD-induced responses were observed (0.0, n=10) (****P <0.0001) (**Fig. 3A,C**).

**Fig. 3.**
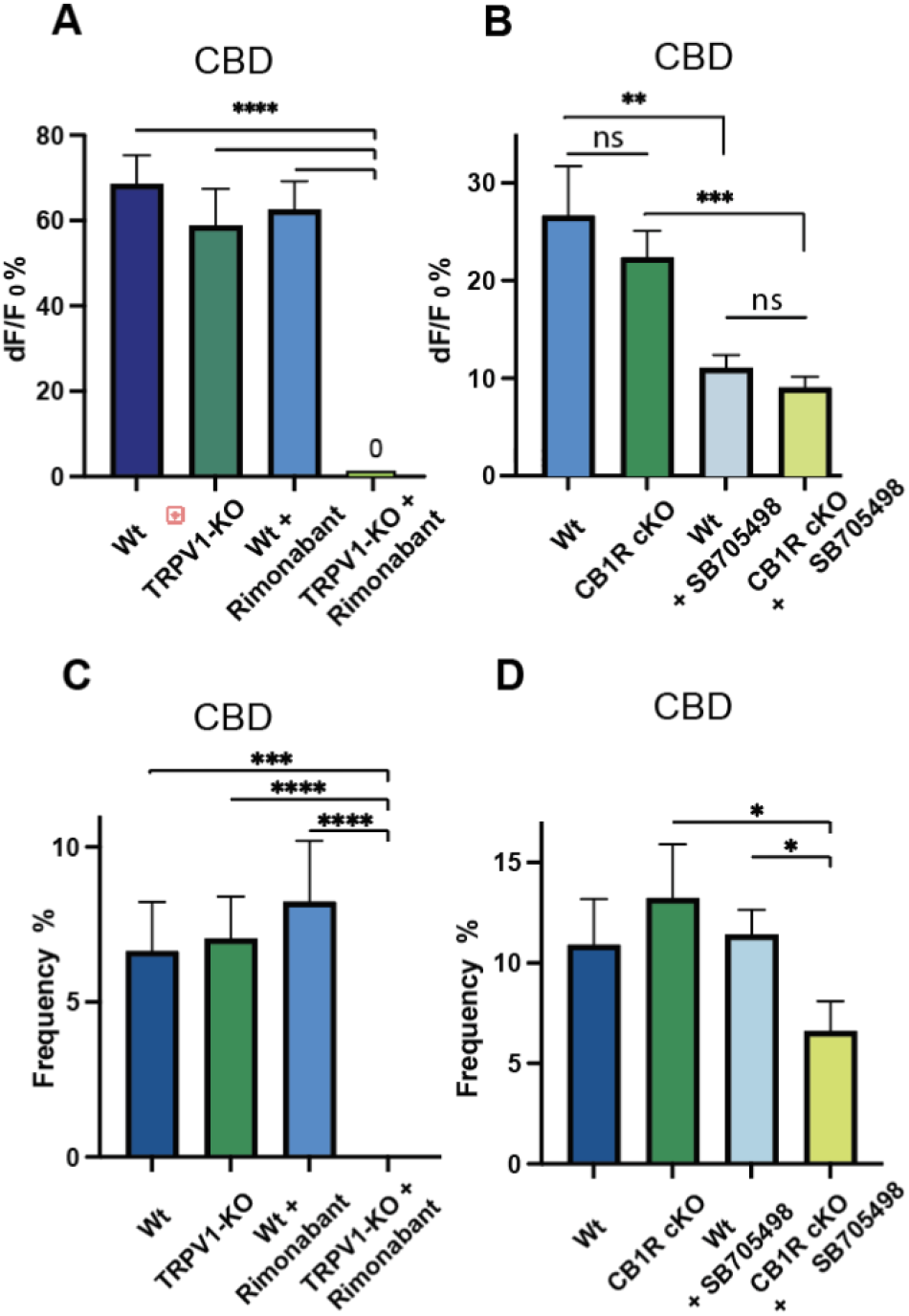
CBD-induced activation of DRG neurons is dependent on co-expression of TRPV1 and CB_1_R. CBD (50μM) - induced DRG neuron [Ca^2+^]_i_ responses (**A**) were unchanged under conditions of genetic TRPV1 knockout (KO) or blockade by the CB_1_R antagonist, Rimonabant (1μM). However, a combination of TRPV1 KO plus Rimonabant completely eliminated CBD-induced responses (**** P< 0.0001). (**B**) Conditional knockout of CB1R (CB_1_R/Nav1.8 (cKD) alone failed to block CBD-induced responses (ns = not significant) but were decreased ∼50% with the TRPV1 antagonist (SB705498) (10μM) alone (** P = 0.004) or in combination with CB_1_R cKO (*** P = 0.0001). (**C**) The frequency (%) of CBD-induced responses was not changed by individual TRPV1 KO or CB_1_R antagonist treatment, but no CBD responses were observed when genetic and pharmacologic blockade were combined. (**D**) Following conditional knockout of CB_1_R or by a TRPV1 antagonist, the frequency of CBD-induced responses remained unchanged. In contrast, the combination of genetic (CB_1_R) and pharmacologic (TRPV1) blockade decreased the frequency of CBD-responding neurons by 40-50% (* P = 0.041). Mean values +/-SEM representing values from ≥ 3 independent trials. Differences were determined by ANOVA with Sidak’s corrections for multiple comparison (A, B), Welch’s correction (C, D) (Graph Pad). Only DRG neurons responsive to KCl at the conclusion of each experiment were included.

Taking a complimentary approach, we repeated the experimental strategy with DRG neurons derived from Wt versus CB_1_R / NaV1.8 cKD mice in the absence or presence of the TRPV1 antagonist SB705498 (TRPV1 blockade validated in **Fig. S3**) [22]. As shown in **Fig 3B**, CBD-induced neuronal responses derived from Wt mice (26.6 +/-5.0, n=21) did not differ from CB_1_R cKD mice (22.4 +/-2.7, n=27). However, the application of CBD in the presence of the TRPV1 antagonist SB705498 showed a 58% decrease (11.0 +/-1.3, n=27) (**P = 0.004). The combination of TRPV1 antagonist SB705498 to neurons from CB_1_R cKD mice directed no additional decrease in response magnitude (59%, 9.0 +/-1.1, n=19; *** P = 0.0001). However, the response frequencies (**Fig. 3D**) decreased under the combination of TRPV1 antagonist plus CB_1_R cKD (6.6 +/-1.4 %, n= 19; *P = 0.041).

### CBN-induced activation of DRG neurons is independent of TRPV1 and CB_1_R expression

There is a limited understanding of CBN’s receptor-mediated antinociceptive properties or if CBN shares similar activation properties with other phytocannabinoids such as CBD [23-25]. For example, CBN was proposed to activate and potentially desensitize smaller-diameter capsaicin-responsive sensory neurons independent of CB_1_-CB_2_-receptors [26]. Given that CBN-induced activation of DRG neurons differed from CBD based on response magnitudes (**Fig. 1E**), sensory neuronal size (larger) (**Fig. 1G**) and dose-response profiles (**Fig. 2A,C**), we investigated the dependence of CBN activation on TRPV1 and CB_1_R in DRG neurons. In contrast to CBD, we found no differences in CBN’s response magnitudes or frequencies under conditions of TRPV1 KO plus a CB_1_R antagonist (**Fig. 4A,C**). We also examined CBN-induced responses in DRG neurons from Wt versus CB_1_R / Adv Cre cKD mice and again found no difference in CBN response magnitudes or response frequency (**Fig. 4B,D**). Therefore, in contrast to CBD, CBN showed no dependence on TRPV1 / CB_1_R-mediated activation of sensory neurons.

**Fig. 4.**
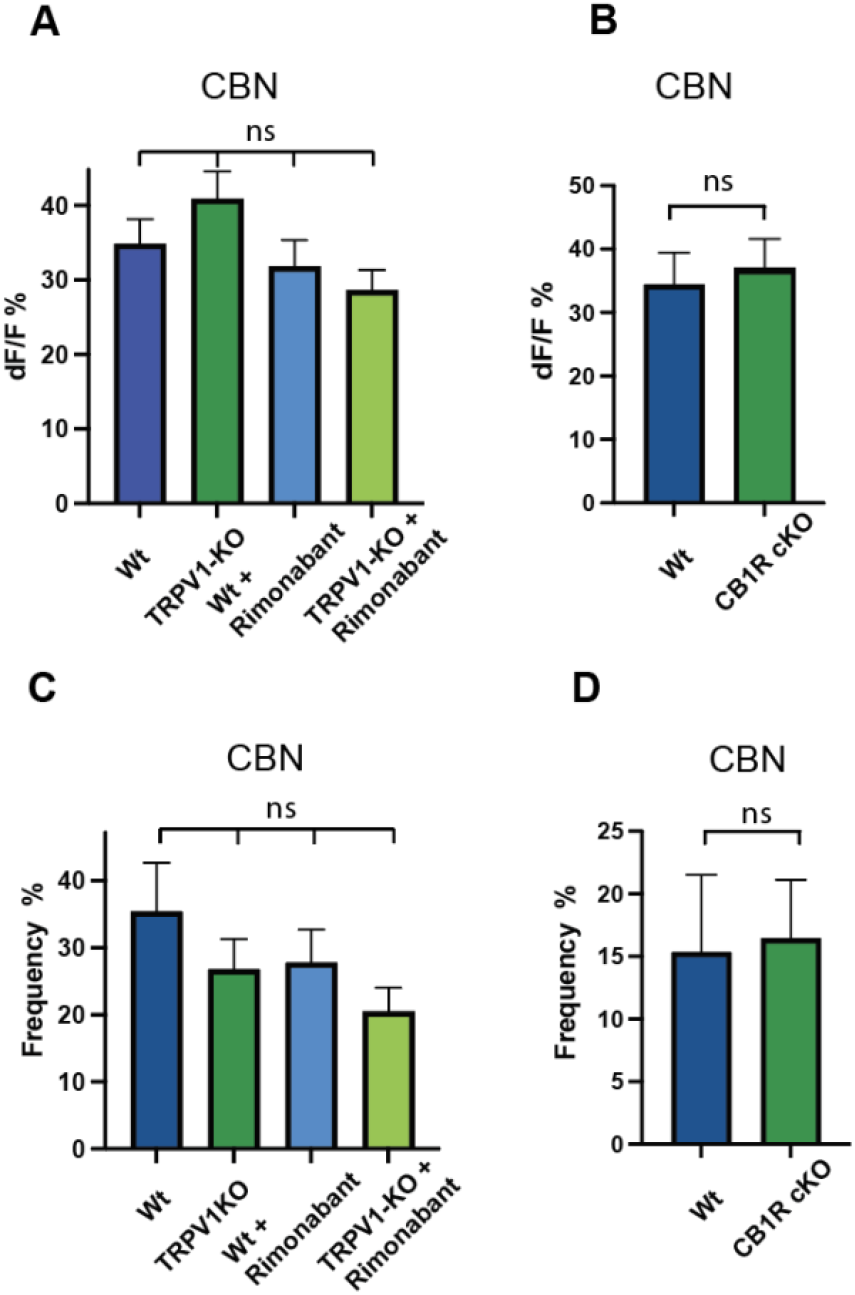
CBN-induced activation of DRG neurons is independent of TRPV1 and/or CB_1_R expression. CBN (50μM) - induced DRG neuron [Ca^2+^]_i_ responses (dF/F%) were unchanged under conditions of (**A**) genetic TRPV1 knockout (KO), blockade by the CB_1_R antagonist, Rimonabant (1μM) or their combination. (**B**) CBN-induced responses did not change under conditions of conditional CB_1_R/Adv KO. The frequency of CBN-induced responses also did not differ under conditions of (**C)** TRPV1 KO, CB_1_R antagonist or their combination. (**D**) The frequency of CBN-induced responses did not differ under conditions of CB_1_R cKO. (ns = not significant). Mean values +/-SEM representing values from ≥ 3 independent trials. Differences were determined by ANOVA with Sidak’s corrections for multiple comparison (**A, C**) or two-tailed t-test (**B, D**) (Graph Pad).

## Discussion

Increasingly, minor cannabinoids such as CBD, CBG, CBN and CBC are being consumed to abate a wide range of undesired maladies including pain and inflammation given their lower profile of unwanted side effects. Moreover, minor cannabinoids CBG, CBC and CBN also give rise to bioactive metabolites via human liver microsomes and may direct additional anti-nociceptive and/or anti-inflammatory properties [27-29]. Despite these advances, our understanding of the mechanisms underlying their proposed benefits is fragmented and often confused with the action of Δ^9^THC. While the receptor-mediated action of minor cannabinoids has been described through their study on individually expressed receptors / channels [9, 10], how those findings translate to the sensory neuronal ecosystem that drives peripheral pain transduction are poorly defined. In this study, we identified differences in minor cannabinoid response magnitudes, size-dependent activation preferences and differences in dose-response relationships in DRG neurons. In the case of CBD and CBN, we also determined to what degree DRG neuron expression of TRPV1 and CB_1_R was necessary for neuronal activation.

### Minor cannabinoids directed different dose-dependent profiles of sensory neuronal activation

Using a concentration of minor cannabinoids (50μM) reported by others to activate and in some cases, desensitize nociceptive transduction [16], we observed that maximal responses induced by CBD, CBG or CBC did not differ from each other but individually were greater than those induced by CBN (**Fig. 1E**). Given that differences in sensory neuronal size has been linked to nociceptive phenotype such as sensitivity to noxious stimuli [18], we next measured the size distribution of DRG neurons activated at 50μM. Surprisingly, all minor cannabinoids tested activated DRG neurons with an equivalent mean size (μm^2^) that differed from the mean value of smaller DRG neurons that were activated by the TRPV1 agonist - capsaicin (**Fig. 1F**) [17]. Notably, when the size distribution of each activating cannabinoid was plotted and compared, differences emerged. For example, CBD and CBG activated the greatest size range, including overlap with smaller capsaicin – responsive neurons. In contrast, CBN and CBC primarily activated a larger-size subpopulation of DRG neurons (**Fig. 1G**). Curiously, CBD and CBG are not known to evoke a painful burning sensation as observed with capsaicin when applied topically. This disparity has been proposed to be related to a difference between capsaicin and CBD in inducing changes in TRPV1 channel pore dilation resulting in higher levels of permeability of Na^+^ and Ca^++^ flux induced by capsaicin [10].

Taken together, CBD and CBG directed dose-dependent increases in calcium responses and activated overlapping neuronal size populations, including capsaicin-sensitive neurons. However, they differed from CBN which activated predominantely *larger* DRG neurons. Importantly, CBN is distinguished from CBD, CBG and CBC where progressively higher concentrations of CBN directed an inverted U-shaped dose - response profile. While we could not identify a prior report of CBN-induced inverted U-shaped dose response curve in sensory neurons, such an inverted U-shaped dose relationship was reported for CBD in directing an antianxiety effect in mice that was later linked to the 5-HT_1A_ receptor [30, 31]. Finally, we observed CBC-induced response properties that included a plateau of 50% from 0.1 - 10 μM followed by higher dose-dependent responses (**Fig. 2D**). Such a biphasic dose-response relationship has been described previously for other receptor systems [32]. Overall, our findings demonstrate that CBD, CBG, CBN and CBC each engender unique properties of size and/or dose-dependent sensory neuronal activation.

### CBD-induced activation of sensory neurons is dependent on co-expression of TRPV1 and CB_1_R

The consequence of TRPV1 and CB_1_R co-expression in sensory neurons has been a focus of study in understanding the peripheral analgesic action of cannabinoids. CBD activates / desensitizes TRPV1 when expressed individually in non-neuronal cell lines [9] and the constitutive activity of CB_1_R in DRG neurons is proposed to maintain TRPV1 in a sensitized state to noxious stimuli [33]. CB_1_R was shown to mediate the peripheral analgesic action of the synthetic THC analog - WIN 55212,22 as demonstrated by a decrease in WIN55212,22’s antinociceptive action in CB_1_R knockdown mice – although the role of CBD was not tested [20].Therefore, we compared CBD-induced responses in primary DRG neurons under genetic knockdown and/or pharmacologic blockade of TRPV1 and/or CB_1_R. Surprisingly, neither genetic knockdown of TRPV1 or pharmacologic blockade (Rimonabant) of CB_1_R alone, decreased CBD-induced activation. However, when both TRPV1 KO plus the CB_1_R antagonist were combined, no measurable CBD-induced responses were observed (**Fig. 3A**). These findings support the hypothesis that CBD-induced activation of DRG neurons is primarily dependent on positive cooperativity between TRPV1 and CB_1_R. Similarly, a model of positive cooperativity was proposed for anandamide and capsaicin-induced activation of DRG neurons via the co-expression of TRPV1 and CB_1_R within a critical cellular distance [21].

Taking a complimentary approach, genetic knockdown of CB_1_R and pharmacologic blockade of TRPV1, resulted in a more complex finding. While conditional knockdown of CB_1_R alone failed to effect CBD-induced activation, application of the TRPV1 antagonist (SB705498) alone decreased CBD-induced response magnitudes. This was unexpected given genetic knockout of TRPV1 alone (**Fig. 3A**) failed to block CBD-induced responses. Nevertheless, the combination of CB_1_R cKD plus TRPV1 antagonist (SB705498) decreased response magnitudes and additionally, the frequency of CBD-induced responses (**Fig. 3D**). We speculate that a lack of concordance of our findings with TRPV1 KO mice and application of the TRPV1 antagonist SB705498 may depend on the ability of the SB705498 to block CBD at the CB_1_R receptor. This is plausible given a report that SB705498 decreased CBD-induced activation of DRG neurons [34].

Another candidate TRP channel reported to mediate CBD-induced activation of sensory neurons is TRPV2 [35]. Given TRPV1 and TRPV2 can form heteromeric complexes in DRG neurons, the contribution of TRPV2 in CBD-induced activation is plausible [36]. Nevertheless, together with published accounts, we propose that CBD is unique in its property of sensory neuronal activation directing the majority of its CBD-induced activation via the co-expression of TRPV1 and CB_1_R in DRG neurons.

### CBN-induced activation of sensory neurons is independent of TRPV1 and CB_1_R

Cannabinol (CBN) a oxidative degradation product of Δ^9^THC is unique in its physiologic and pharmacologic properties [25]. Most notable is that CBN lacks the majority of psychomimetic effects associated with Δ^9^THC while retaining certain antinociceptive properties [24, 37]. However, the mechanism in which CBN directs antinociceptive action via sensory neurons is not understood. In contrast to our findings for CBD, when CBN was applied to cultured DRG neurons under conditions of TRPV1 KO plus the CB_1_R antagonist Rimonabant, no difference in response magnitudes or frequency was observed (**Fig. 4**). When an identical experiment (Wt versus TRPV1-KO, Rimonabant or TRPV1-KO plus Rimonabant) was repeated with DRG neurons from female mice (**Fig. S4**), we also observed no difference in the magnitude of CBN-induced responses (dF/F%). Therefore, despite reports of CBN-induced activation of TRPV1 in transfected HEK cells [10], and in contrast to our findings for CBD, CBN-induced activation of DRG neurons appeared independent of TRPV1 / CB_1_R as summarized in **Fig. 5**. These findings are concordant with earlier studies describing CBN-induced, CGRP mediated relaxation of arterial vessels that were independent of TRPV1, based on pharmacologic blockade [26].

**Fig. 5.**
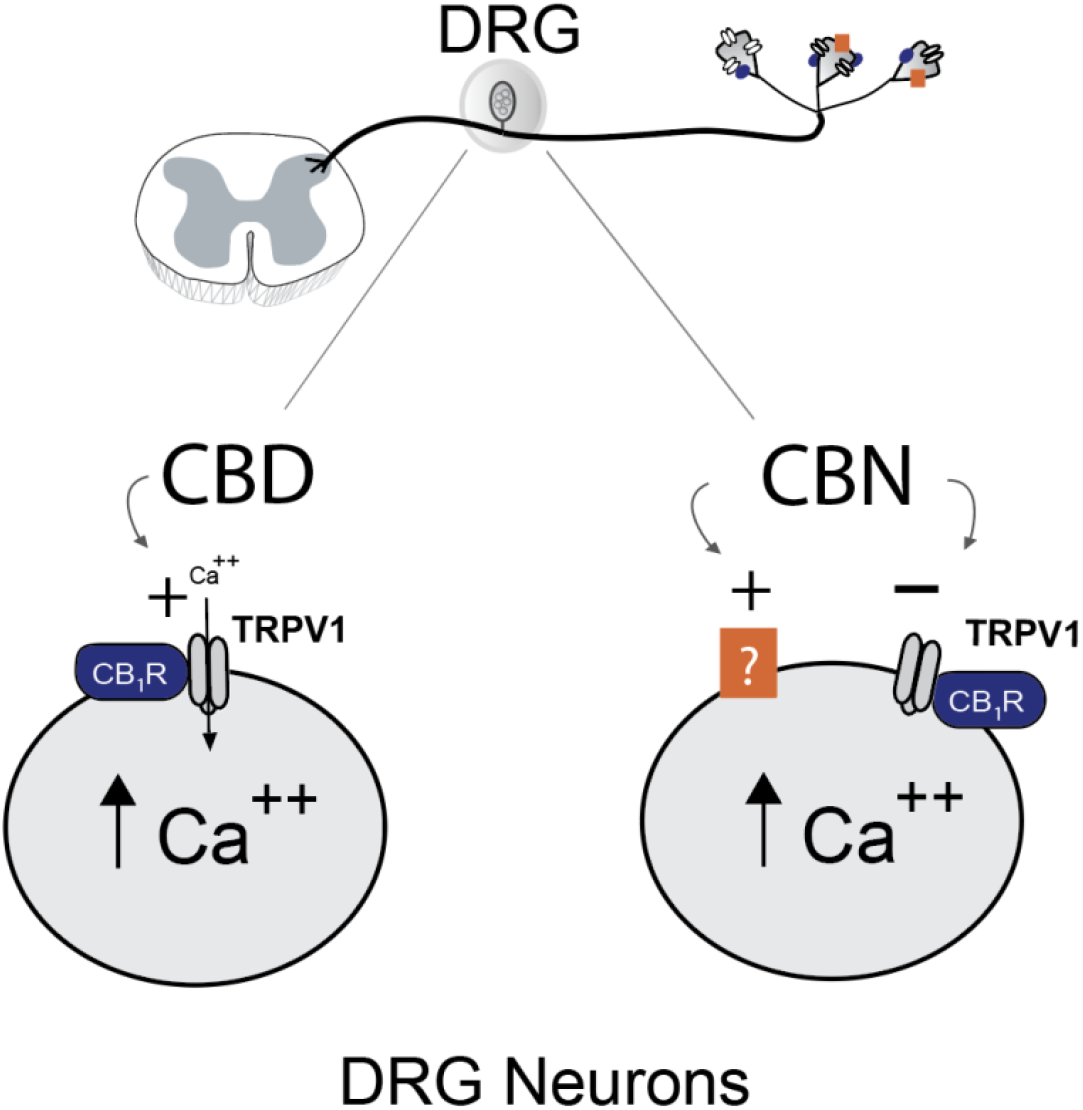
CBD and CBN differ in their receptor-mediated activation of DRG neurons. Summary illustration denoting the activation (increased [Ca^2+^]_i_) of hypothetical sensory neurons by CBD and CBN. **CBD**-induced activation of sensory neurons is proposed to be dependent on the co-expression of the multimodal pain transducing channel TRPV1 and cannabinoid receptor CB_1_R. Genetic loss of TRPV1, CB_1_R, or pharmacologic blockade of CB_1_R did not inhibit CBD-induced responses. However, the combination of TRPV1 KO plus pharmacologic blockade of CB_1_R eliminated all CBD-induced responses. While CBD is capable of activating a broad size range of sensory neurons as modeled by cultured DRG neurons, CBD directed a linear dose-response profile consistent with a mechanism of cooperativity. **CBN**-induced activation of larger sensory neurons directed an inverted U-shaped dose-response profile. Therefore, activation by CBN is proposed to be independent of TRPV1 and/or CB_1_R given that genetic loss of TRPV1 and CB_1_R, as well as pharmacologic blockade of CB_1_R, individually or in combination, with TRPV1 knockout did not inhibit CBN-induced responses. Identifying the channel / receptor(s) that mediate CBN-induced activation of sensory neurons requires further study.

In the search for alternative receptors mediating peripheral cannabinoid analgesia, orphan receptor GPR55 was investigated and found to be activated by Δ^9^THC and anandamide [38]. However, GPR55 was later identified as the lysophosphatidylinositol 1 receptor [39, 40] and found not to be required for inflammatory and neuropathic nociception [41]. While efforts are ongoing to link orphan receptors to cannabinoid analgesic action [42], the inhibitory action of minor cannabinoid CBN on sodium channels (NaV1.8) expressed on nociceptors holds promise as a plausible antinociceptive mechanism [43, 44]. Nevertheless, it remains to be determined what receptor / channel(s) mediate CBN-induced activation and subsequent desensitization of DRG neurons.

### Limitations

In contrast to dose-response profiles derived from minor cannabinoid activation of an individual channel / receptor (eg. TRP channels) expressed in an established cell line, we undertook a comparison of response properties across a phenotypically diverse population of sensory neurons. Differences in minor cannabinoid dose-response curves, sensory neuronal size preference and dependence (CBD) or independence (CBN) from TRPV1 confounded application and comparison of EC_50_ values across experimental strategies. As such, comparative studies utilized peak DRG neuron calcium responses induced by cannabinoids at 50 μM under control, pharmacologic blockade or from neurons derived from genetically modified mice. Therefore, these comparative studies do not preclude additional outcomes at concentrations below or above 50 μM.

### Conclusion

We observed that minor cannabinoids CBD, CBG, CBN and CBC differed in their activation of DRG neurons. Notably, CBD displayed a linear dose-response relationship directed by DRG neurons that supported a model of positive cooperativity and striking requirement for co-expression of TRPV1 and CB_1_R. In contrast, CBN-induced activation was distinguished by its preference for larger DRG neurons, an inverted U-shaped dose-response profile and activation of DRG neurons that was independent of TRPV1. Therefore, these four minor cannabinoids directed unique activation properties across a diverse population of sensory neurons revealing an opportunity to investigate their potential antinociceptive entourage effects.

## Acknowledgments

We thank Drs. Che-Chung Yeh UCSF for his expertise in breeding and managing CB_1_R / Nav1.8 cKD mice and Dr. Jarret Weinrich for insightful discussion of the manuscript (Department of Anesthesia and Perioperative Care). This work was supported by research grants from NIH/NCCIH R01 AT010757, California Department of Cannabis Control (DCC) and the UCSF Dept of Anesthesia and Perioperative Care.

## Notes

### Competing Interest Statement

The authors have declared no competing interest.

