## Supplementary material for "Minor Cannabinoids CBD, CBG, CBN and CBC differentially modulate sensory neuron activation": Fig. S1

**Associated Data Supplementary Figures**

### Minor Cannabinoids CBD, CBG, CBN and CBC differ in their properties of sensory neuronal activation

**Running Title: Minor cannabinoids and sensory neuron activation**

#### CBD

| Time | % of the initial concentration | % Total Impurities |
|---|---|---|
| Initial | 100% | 0.2% |
| Day 7 | 100% | 0.0% |
| Day 14 | 100% | 0.0% |
| 1 Month | 101% | 0.0% |
| 8 Months* | 104% | 0.4% |

#### CBG

| Time | % of the initial concentration | % Total Impurities |
|---|---|---|
| Initial | 100% | 0.7% |
| Day 7 | 101% | 0.9% |
| Day 14 | 101% | 0.9% |
| 1 Month | 101% | 0.3% |
| 8 Months* | 102% | 0.2% |

#### CBN

| Time | % of the initial concentration | % Total Impurities |
|---|---|---|
| Initial | 100% | 1.0% |
| Day 7 | 100% | 1.1% |
| Day 14 | 100% | 0.8% |
| 1 Month | 101% | 0.7% |
| 8 Months* | 107% | 1.0% |

#### CBC

| Time | % of the initial concentration | % Total Impurities |
|---|---|---|
| Initial | 100% | 2.1% |
| Day 7 | 100% | 2.0% |
| Day 14 | 100% | 2.3% |
| 1 Month | 100% | 0.3% |
| 8 Months* | 108% | 0.8% |

**Fig. S1.** Stability and purity of minor cannabinoid stock solutions in ethanol at -20°C over a period of eight months. (*) An increase in concentration was attributed to evaporation of ethanol.

| Cannabinoid | % of Target Concentration as Prepared | Initial % of Target Concentration in Apparatus | % of Target Concentration in Apparatus at 1 hr |
|---|---|---|---|
| CBD | 86% | 82% | 71% |
| CBN | 79% | 75% | 71% |
| CBG | 74% | 62% | 64% |
| CBC | 96% | 91% | 94% |

**Fig. S2A. Changes in the concentration of minor cannabinoids in 50 μM solutions prepared for calcium imaging with DMSO.** A trial to prepare CBD in aqueous assay buffer with 0.2% DMSO yielded a solution that was below the target concentration and the solution continued to lose CBD over 1hr at room temperature (RT). Agitation and/or sonication increased the rate of loss as did increasing the DMSO concentration to 1%.

| Day | [μM] |
|---|---|
| 0 | 31 |
| 9 | 4.5 |
| 14 | 3.2 |
| 16 | 2.7 |
| 23 | 2.4 |

**Fig. S2B. Stability of CBD 50 μM in assay buffer with DMSO (at RT)** CBD concentration declined over 23 days in a 1.8 mL glass HPLC vial at RT.

| Condition | [μM] |
|---|---|
| Glass syringe @ 0 hr | 45 |
| Glass syringe @ 1 hr | 42 |
| Sonication @ 0 | 43 |
| Plastic tubing at 0 min | 38 |
| Plastic tubing at 1 hr | 3 |

**Fig. S2C. Preparation of CBD 50 μM in assay buffer with ethanol.** A CBD solution was prepared by adding 100 μL of a 1mg/ml stock CBD solution in ethanol directly to the assay buffer at RT.

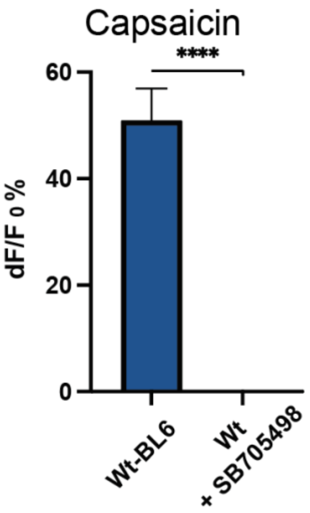


**Fig. S3 Validation of TRPV1 antagonist SB705498** The TRPV1 antagonist SB705498 (10μM) completely inhibited capsaicin (1μM) - induced DRG neuron responses. 3 independent trials, n= 25 per group (**** $p<0.0001$). Statistical comparison: unpaired t-test. Mean +/- SEM. DRG, dorsal root ganglion, Wt: Wildtype.

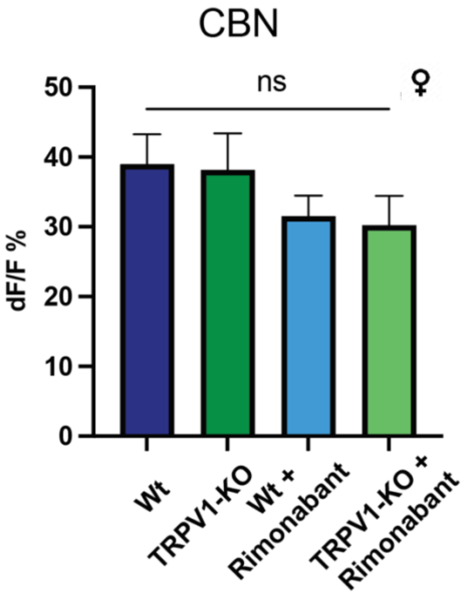


**Fig. S4 Female mouse DRG neuron responses to CBN.** CBN (50μM) - induced response magnitudes in female DRG neurons were not significantly changed under conditions of TRPV1 knockout (KO), $CB_1R$ antagonist Rimonabant (1μM) alone or in combination with TRPV1 KO. n = 53, 31, 50, 31; (p = ns) ANOVA with Tukey's test for multiple comparisons, 3 independent trials. Mean +/- SEM. DRG, dorsal root ganglion. ns: not significant.
